# Bacterial profiles and their antimicrobial susceptibility pattern of Isolates from inanimate hospital environments at Tikur Anbessa Specialized Teaching Hospital, Addis Ababa, Ethiopia

**DOI:** 10.1101/2020.09.25.312983

**Authors:** Shemse Sebre, Woldaregay Erku, Aminu Seman, Tewachw Awoke, Zelalem Desalegn, Wude Mihret, Adane Mihret, Tamrat Abebe

**Affiliations:** Department of Microbiology, Immunology and Parasitology, School of Medicine, College of Health Sciences, Addis Ababa University, Addis Ababa, Ethiopia; Armauer Hansen Research Institute; Addis Ababa, Ethiopia; Department of Medical Laboratory Sciences, College of Health Sciences, Bahir Dar University, Bahir Dar, Ethiopia

**Keywords:** Antimicrobial susceptibility pattern, Operation theaters, Inanimate Hospital environments, Intensive care unit, Bacteria

## Abstract

Microbial contamination of hospital environment plays an important role in the spread of health care-associated infections (HCAIs). This study was conducted to determine bacterial contamination, bacterial profiles and antimicrobial susceptibility pattern of bacterial isolates from environmental surfaces and medical equipment. A cross-sectional study was conducted at Tikur Anbessa Specialized Hospital (TASH) from June to September, 2018. A total of 164 inanimate surfaces located at intensive care units (ICUs) and operation theaters (OTs) were swabbed. All isolates were identified by using routine bacterial culture, Gram staining and a panel of biochemical tests. For each identified bacteria, antibiogram profiles were determined by the Kirby Bauer disk diffusion method according to the guidelines of the Clinical and Laboratory Standards Institute (CLSI). Out of the 164 swabbed samples, 141 (86%) were positive for bacterial growth. The predominant bacteria identified from OTs and ICUs were *S. aureus* (23% vs 11.5%), *Acinetobacter* spp (3.8% vs 17.5%) and Coagulase negative *Staphylococcus* (CONS) (12.6% vs 2.7%) respectively. Linens were the most contaminated materials among items studied at the hospital (14.8%). The proportions of resistance among Gram-positive bacteria (GPB) were high for penicillin (92.8%), cefoxitin (83.5%) and erythromycin (54.1%). However, the most effective antibiotics were clindamycin with only 10.4% and 16.5% resistance rates, respectively. The antimicrobial susceptibility profiles of Gram-negative bacteria (GNB) revealed that the most effective antibiotics were amikacin, ciprofloxacin, and gentamicin with resistance rate of 25%, 37.5%, and 46.3%, respectively. However, the highest resistance was recorded against ampicillin (97.5%), ceftazidime (91.3%), ceftriaxone (91.3%) and aztreonam (90%). The inanimate surfaces near immediate patient environment and commonly touched medical equipment within OTs and ICUs are reservoirs of potential pathogenic bacteria that could predispose critically ill patients to acquire HCAIs. The proportions of antimicrobial resistance profile of the isolates are much higher from studied clean inanimate environments.

## Introduction

Hospital environment represents a new ecological place for medically important nosocomial pathogens, antibiotic-resistant microorganisms and reservoirs of resistance gene, which have been commonly, found on various surfaces within hospitals (e.g. medical equipment, housekeeping surfaces, workplaces and lobby (furniture) [1, 2]. Studies investigating hospital environments reported that pathogens were ubiquitous in all hospital units but the interest was usually focused on intensive care and operation unit, especially due to the vulnerability of patients in these units [3]. There is also high antibiotic usage and invasive procedure from these units [1].

Bacterial cross-contamination plays an important role in health care-associated infections (HCAIs) and resistant strain dissemination [1, 4]. The majority of the HCAIs are believed to be transmitted directly from patient to patient, but increasing evidence demonstrates that also the medical personnel as well as the clinical environment (i.e., surfaces and equipment) often are a source of infections [5]. Hospital design and hygienic practices have been largely directed at controlling nosocomial pathogens and resistant strains contaminating air, hands, equipment, and surfaces [6]. A better understanding of how bacterial cross-contamination occurs can provide the basis for the development of evidence-based preventive measures [4].

Emergence of multi-drug resistant (MDR) strains in a hospital environment; particularly in developing countries, is an increasing problem which is an obstacle for management of HCAIs [7–10]. In Ethiopia, studies reported high prevalence of HCAIs mainly due to MDR pathogens including the country’s largest tertiary referral Hospitals [11–13], which warrants the critical need for a reassessment of the role played by inanimate environment in the transmission of nosocomial infections [6, 14].

Studies on the bacterial contaminations of ward of the hospital environments in Ethiopia reported high bacterial load and multidrug resistant (MDR) strains [9, 10, 15, 16]. However, few data exist on the bacterial contamination of the hospital environment in the studied hospital. Therefore, the aim of this study were to determine bacterial contamination, detect potential pathogenic bacteria and to determine the antimicrobial susceptibility patterns from inanimate hospital environments in the environments of Operation Theaters (OTs) and Intensive Care Units (ICUs) at Tikur Anbessa Specialized Teaching Hospital in Addis Ababa, Ethiopia.

## Materials and Methods

### Study setting, Study period and Sampling locations

A cross-sectional study was conducted at Tikur Anbessa Specialized Hospital (TASH), Addis Ababa, Ethiopia from June to September, 2018. TASH is a tertiary hospital and major referral center for other hospitals in Ethiopia. TASH has 800 beds and provides care for approximately 370,000-400,000 patients per year. The samples were collected from four intensive care units including Surgical, Pediatric, Medical and Medical-Surgical units. A total of seven operating theaters were examined including Emergency, Neurology, Endo-Renal, Obstetrics and gynaecology, Pediatrics, Cardio-Vascular and Gastro intestinal tract (GIT) units.

### Surfaces sampling

The detection of bacteria in ICUs and OTs were performed by using the swab method from surfaces and medical devices. All samples were collected every morning after cleaning of the hospital environment was completed. Moreover, samples in OTs were collected before start of operations. Sampling sites around a bed in each ICUs and OTs were chosen based on the frequency with which the surfaces were touched. Sterile swabs were moistened in Brain Heart Infusion (BHI) and then, were used to swab (i) commonly touched medical equipment including beds, monitors, OR-light, linens, ventilators, oxygen supply, anesthesia machine, suction buttons and Laparoscopy (ii) workstation, including keyboards, computer mice; (iii) environments including floors, wall and corridors; (iv) Lobby (furniture) including chair, table, lockers and trowels; (v) Sinks; (vi) hospital textiles including bed linen based on methods described previously [17–20].

### Microbiology Analysis

Each swab sample was pre-enriched in sterile BHI and incubated at 37°C for 24 hours. A loop full of the turbid broth was then sub-cultured on blood agar (Oxoid, UK), Mannitol salt agar (MSA), MacConkey agar and Chromagar TM Strep B base plates (Chromagar microbiology, France). Differential and selective characteristics for each agar medium were recorded for the initial screening of suspected potential pathogens. Furthermore, specific colony color (mauve color) on Chromagar TM Strep B was considered for Group B *Streptococci* (GBS) while yellow colony color on MSA was considered for *S. aureus*.

Gram-negative bacteria were further identified by Gram stain and standard biochemical tests like Triple Sugar Iron Agar (TSI), urea, citrate, Sulfide Indole Motility (SIM) medium, growth in Lysine Iron Agar (LIA), Mannitol, malonate, and oxidase test. On the other hand, Gram-positive bacteria were further identified by Gram stain, optochin, bacitracin, CAMP test and different biochemical tests such as catalase, coagulase, bile esculin and salt tolerance test described based on hand book of Clinical Microbiology Procedures [21].

### Antimicrobial susceptibility testing

Antimicrobial susceptibility testing of the isolates were performed using 21 antibiotics (Oxoid, UK) based on the Kirby-Bauer disk diffusion method on Mueller-Hinton agar (MHA) (Oxoid, UK) and Mueller-Hinton with blood agar (Oxoid, UK) for *Streptococci* spp and *Enterococcus* spp [22]. An inoculum for each isolate was prepared by emulsifying colonies from an overnight pure culture in sterile normal saline (0.85%) in test tubes with the turbidity adjusted to 0.5 McFarland standards. The bacterial suspension was uniformly streaked on MHA plates using sterile swabs and left for 3 minutes prior to introduction of the antibiotics.

For Gram-negative bacteria the following antibiotics were used (in μg/disk): ampicillin (10), amoxicillin and clavulanic acid (10/10), ceftriaxone (30), cefotaxime (30), ceftazidime (30), amikacin (30), gentamicin (10), ciprofloxacin (5), sulfamethoxazole-trimethoprim (1.25/23.75), cefoxitin (30), cefuroxime (30), cefepime (30), piperacillin-tazobactam (100/10), meropenem (10) and aztreonam (30) based on Clinical Laboratory Standards Institute (CLSI) [22].

On the other hand, for Gram-positive bacteria antibiotics (in μg/disk) selected for susceptibility testing included penicillin (10 units), gentamicin (10), erythromycin (15), ciprofloxacin (5), doxycycline (30), vancomycin (30), cefoxitin (30), sulfamethoxazole-trimethoprim (1.25/23.75), clindamycin (2) and chloramphenicol (30). The plates were incubated at 35 ^0^C for 24 h, and the diameters of zone of inhibition were measured with Vernier caliper and results were reported as susceptible (S), intermediate (I), or resistant (R), according to CLSI guidelines [22].

### Quality Assurance

To ensure the quality of the result from different assays, internal quality assurance systems was in place for all laboratory procedures and double checking of the result was done. All the methods to be used were validated as fit for the purpose before use in the study. Standard operating procedures (SOPs) were used for specific purpose for all laboratory procedures. Quality control strains of *Enterococcus faecalis* ATCC® 29212, *S. aureus* ATCC® 25923, *E. coli* ATCC® 2592, *K. pneumoniae* ATCC®1705 and *K. pneumoniae* ATCC®1706 were used to confirm the result of antibiotics, media and to assess the quality of the general laboratory procedure [22].

### Statistical analysis

Data analysis was performed using Stata version 14 software program (Stata Corporation, Lakeway Drive, College Station, Texas), and descriptive statistics (percentages or frequency) was calculated. A difference was considered statistically significant for P-value ≤ 0.05.

### Ethics approval

The study protocol was approved by the Department of Microbiology, Immunology and Parasitology Research Ethics Review Committee (DRERC), College of Health Sciences, Addis Ababa University (Ref. no. DRERC/17/18/02-G). Prior to sample collection, written approval was obtained from administrative unit of Tikur Anbessa Specialized Hospital.

## Results

### Culture Results

During the four months study, a total of 164 environmental swabs were collected in the studied OTs (n=99) and ICUs (n=65) of the hospital. Of these swab samples, 141(86%) were positive for bacterial growth, from which a total of 183 bacterial isolates were identified. Multi-bacterial contamination was detected in 26.8% of the samples, mainly found on the surfaces of ventilators, bed and linens.

### Frequency of bacterial etiologies

Out of the 183 bacterial isolates, 103(56.3%) were Gram-positive bacteria (GPB) and the rest Gram-negative bacteria (GNB). Among the GPB *S. aureus* (34.4%), CONS (15.3%) and *Bacillus* spp (3.3%) were the dominant isolates. Among the GNB *Acinetobacter* spp (21.3%), *Pseudomonas* spp (7.7%) and *E. coli* (4.9%) were the dominant isolates. Overall, *S. aureus* was the most frequently isolated bacteria (34.4%) followed by *Acinetobacter* spp (21.3%) and CONS (15.3%) (Table 1).

**Table 1:** The frequency of isolated bacteria at TASH, 2018

| Isolates | N (%) |
| --- | --- |
| **Gram-negative** | **80(43.7)** |
| *Acinetobacter* spp | 39(21.3) |
| *Pseudomonas* spp | 14(7.7) |
| *E. coli* | 9(4.9) |
| *Serratia* spp | 4(2.2) |
| *Klebsiella pneumoniae* | 6(3.3) |
| *Klebsiella oxytoca* | 4(2.2) |
| Others* | 4(2.2) |
| **Gram-positive** | **103(56.3)** |
| *S. aureus* | 63(34.4) |
| CONS | 28(15.3) |
| *Bacillus* spp | 6(3.3) |
| *Streptococcus agalactiae* | 3(1.6) |
| *Enterococcus* spp | 3(1.6) |

\*Others: (*Enterobacter* spp, *Shigella* spp, *Klebsiella rhinoscleromatis*)

### Distribution of bacterial isolates between ICUs and OTs

Most of the potential bacterial pathogens were isolated from Intensive care units (ICUs), 50.3% (92/183). Significant differences between Gram-positive and Gram-negative bacteria were observed between wards in OTs (39.9% vs 9.8%) and ICUs (16.4% vs 33.9%) respectively (p=0.000). The ICUs were mainly contaminated with GNB, 67.4% (62/92), of which the predominant ones being *Acinetobacter* spp accounting for 34.8% (32/92) followed by *S. aureus* with 22.8 % (21/92) isolation rate. Most of the bacteria in ICUs were isolated from Medical-Surgical (16.4%, 30/183) ward. The major pathogens in this ICU were *S. aureus* from GPB and *Acinetobacter* spp from GNB, each with isolation rate of (33.3%, 10/16). The Operation Theaters (OTs) were mainly contaminated by GPB, 80.2% (73/91). The major pathogens in the theatre were *S. aureus*, 46.2% (42/91) and CONS, 25.3% (23/91). Endo-Renal theatre was mostly contaminated with *S. aureus* with rate as high as 31.3% (5/16) (Table 2).

**Table 2:** Distribution of potential pathogenic bacteria between ICUs and OTs at TASH, 2018

| Bacteria | ICUs (N= 92) |  |  |  | OTs (N=91) |  |  |  |  |  |  |
| --- | --- | --- | --- | --- | --- | --- | --- | --- | --- | --- | --- |
|  | Surgical<br>n (%) | Pediatric<br>n (%) | Medical<br>n (%) | Medical-Surgical<br>n (%) | Emergency<br>n (%) | Neurology<br>n (%) | Endo-Renal<br>n (%) | Gyn-obs<br>n (%) | Pediatric<br>n (%) | Cardo-Vs<br>n (%) | GIT<br>n (%) |
| *S. aureus* | 4(25) | 4(16.7) | 3(13.6) | 10(33.3) | 8(66.7) | 7(53.8) | 5(31.3) | 6(60) | 4(33.3) | 6(42.9) | 6(42.9) |
| CONS | 1(6.3) | 0(0) | 3(13.6) | 1(3.3) | 4(33.3) | 2(15.4) | 0(0) | 3(30) | 6(50) | 6(42.9) | 2(14.3) |
| *Bacillus spp* | 0(0) | 0(0) | 0(0) | 0(0) | 0(0) | 2(15.4) | 0(0) | 1(10) | 0(0) | 0(0) | 3(21.4) |
| *Enterococcus spp* | 0(0) | 1(4.2) | 1(4.5) | 0(0) | 0(0) | 0(0) | 0(0) | 0(0) | 1(8.3) | 0(0) | 0(0) |
| GBS | 0(0) | 0(0) | 0(0) | 2(6.7) | 0(0) | 1(7.7) | 0(0) | 0(0) | 0(0) | 0(0) | 0(0) |
| *Acinetobacter spp* | 6 (37.5) | 6 (25) | 10 (45.5) | 10 (33.3) | 0(0) | 0(0) | 4(25) | 0(0) | 1(8.3) | 1(7.1) | 1(7.1) |
| *Pseudomonas spp* | 2(12.5) | 5(20.8) | 2(9.1) | 0(0) | 0(0) | 1(7.7) | 2(12.5) | 0(0) | 0(0) | 0(0) | 2(14.3) |
| *Klebsiella spp* | 1(6.3) | 4(16.7) | 2(9.1) | 2(6.7) | 0(0) | 0(0) | 2(12.5) | 0(0) | 0(0) | 0(0) | 0(0) |
| *E. coli* | 1(6.3) | 4(16.7) | 1(14.5) | 3(10) | 0(0) | 0(0) | 0(0) | 0(0) | 0(0) | 0(0) | 0(0) |
| *Serratia spp* | 1(6.3) | 0(0) | 0(0) | 0(0) | 0(0) | 0(0) | 2(12.5) | 0(0) | 0(0) | 1(7.1) | 0(0) |
| Others * | 0(0) | 0(0) | 0(0) | 2(6.7) | 0(0) | 0(0) | 1(6.3) | 0(0) | 0(0) | 0(0) | 0(0) |
| **Total, N (%)** | 16 (8.7) | 24(13.1) | 22(12) | 30(16.4) | 12(6.6) | 13(7.1) | 16(8.7) | 10(5.5) | 12(6.6) | 14(7.7) | 14(7.7) |

Others \* (*Shigella* spp, *Enterobacter* spp); GIT: Gastro-intestinal tract unit, Cardo-Vs: Cardiovascular unit; Gyn-obs: Gynaecology obstetrics; GBS: Group B Streptococcus (*Streptococcus agalactiae*), CONS: Coagulase negative staphylococci, OTs: Operation theaters; ICUs: Intensive care units

### Distribution of bacterial pathogens over different surfaces

The highest bacterial contaminated samples were taken from bed linens followed by environmental surfaces and bed. Linens were mostly contaminated with *Klebsiella* spp., (54.5%, 6/27), followed by *Acinetobacter* spp., (15.4%, 6/39). Beds were mainly contaminated with *S. aureus* (12.7%, 8/63). Sinks were mainly colonized by *S. aureus* (7.7%, 6/63), *Pseudomonas* spp (7.1%, 1/14) and *Acinetobacter* spp (5.1%, 2/39). *Klebsiella* spp is mainly contaminated ventilators (27.3%, 3/11) (Table 3).

**Table 3:** Distribution of bacteria over different surfaces in ICU and OTs at TASH, Addis Ababa, Ethiopia, 2018

| Sampling points | Bacteria, n (%) |  |  |  |  |  |  |  |  |  |  |
| --- | --- | --- | --- | --- | --- | --- | --- | --- | --- | --- | --- |
|  | *S. aureus* | CONS | *Bacillus spp* | *Enterococcus spp* | GBS | *Acinetobacter spp* | *Pseudomonas spp* | *Klebsiella spp* | *E. coli* | *Serratia spp* | Others <sup>A</sup> |
| Anastasia machine | 5(7.9) | 2(7.1) | 1(16.7) | 0(0) | 0(0) | 1(2.6) | 1(7.1) | 0(0) | 0(0) | 0(0) | 0(0) |
| Bed | 8(12.7) | 2(7.1) | 0(0) | 0(0) | 1(33.3) | 8(20.5) | 1(7.1) | 1(9.1) | 2(22.2) | 0(0) | 0(0) |
| Environmental surface* | 6(9.5) | 6(21.4) | 3(50) | 1(33.3) | 0(0) | 3(7.7) | 1(7.1) | 1(9.1) | 0(0) | 1(25) | 0(0) |
| Monitor | 5(7.9) | 3(10.7) | 0(0) | 0(0) | 1(33.3) | 4(10.3) | 1(7.1) | 0(0) | 1(11.1) | 0(0) | 1(33.3) |
| Sink | 6(9.5) | 0(0) | 0(0) | 0(0) | 0(0) | 2(5.1) | 1(7.1) | 0(0) | 0(0) | 0(0) | 1(33.3) |
| Suction Machine | 7(11.1) | 3(10.7) | 0(0) | 1(33.3) | 0(0) | 5(12.8) | 3(21.4) | 0(0) | 1(11.1) | 0(0) | 0(0) |
| Linens | 5(7.9) | 4(14.3) | 0(0) | 0(0) | 1(33.3) | 6(15.4) | 3(21.4) | 6(54.5) | 2(22.2) | 0(0) | 0(0) |
| Lobby (furniture) | 5(7.9) | 3(10.7) | 0(0) | 0(0) | 0(0) | 5(12.8) | 0(0) | 0(0) | 1(11.1) | 1(25) | 0(0) |
| Ventilator | 1(1.6) | 1(3.6) | 0(0) | 0(0) | 0(0) | 3(7.7) | 1(7.1) | 3(27.3) | 2(22.2) | 0(0) | 0(0) |
| Work station | 6(9.5) | 1(3.6) | 2(33.3) | 0(0) | 0(0) | 2(5.1) | 0(0) | 0(0) | 0(0) | 2(50) | 0(0) |
| Others <sup>B</sup> | 9(14.3) | 3(10.7) | 0(0) | 1(33.3) | 0(0) | 0(0) | 2(14.3) | 0(0) | 0(0) | 0(0) | 1(33.3) |
| **Total** | **n=63** | **n=28** | **n=6** | **n=3** | **n=3** | **n=39** | **n=14** | **n=11** | **n=9** | **n=4** | **n=3** |

Others <sup>A</sup> (*Shigella* spp, *Enterobacter* spp); Others <sup>B</sup> (Laparoscopy, OR-Light, oxygen cylinder, Trowels); \*Environmental surfaces (Door knob, Floor, Corroder and Wall), Group B Streptococcus (*Streptococcus agalactiae*), CONS: Coagulase negative staphylococci.

### Antibiogram profile for Gram-positive isolates

The proportions of antimicrobial resistance among GPB were high for penicillin (92.8%), cefoxitin (83.5%) and erythromycin (54.1%). Low level of resistance was recorded for clindamycin (10.4%) and gentamicin (16.5%). Using cefoxitin disk as a surrogate marker, 54(85.7%) of *Staphylococcus aureus* isolates were defined as MRSA. High resistance level was also recorded to penicillin (93.7%). Vancomycin resistance was demonstrated by 12 (19%) *S. aureus*, 5 (17.9%) CONS and 1(33.3%) *Enterococcus* spp (Table 4).

**Table 4:** Antimicrobial susceptibility pattern of Gram-positive bacteria at TASH, 2018.

| Isolates | Antimicrobial agents N (%) |  |  |  |  |  |  |  |  |  |  |
| --- | --- | --- | --- | --- | --- | --- | --- | --- | --- | --- | --- |
|  | Ptn | GEN | CIP | CHL | SXT | VAN | ERY | DA | DOX | FOX | PEN |
| **CONS** | R | 5(17.9) | 10(32.1) | 6(21.4) | 11(39.3) | 5(17.9) | 16(57.1) | 1(3.6) | 15(53.6) | 22(78.6) | 26(92.9) |
|  | S | 23(82.1) | 18(64.3) | 22(78.6) | 17(60.7) | 23(82.1) | 12(42.9) | 27(96.4) | 13(46.4) | 6(21.4) | 2(7.1) |
| *Enterococcus spp* | R | NT | 1(33.3) | 1(33.3) | NT | 1(33.3) | 2(67.7) | 2(66.7) | 1(33.3) | NT | 2(66.7) |
|  | S | NT | 2(66.7) | 2(67.7) | NT | 2(66.7) | 1(33.3) | 1(33.3) | 2(66.7) | NT | 1(33.3) |
| **GBS** | R | NT | NT | 2(66.7) | NT | 2(66.7) | 3(100) | 0(0) | 1(33.3) | NT | 3(100) |
|  | S | NT | NT | 1(33.3) | NT | 1(33.3) | 0(0) | 3(100) | 2(66.7) | NT | 0(0) |
| *S. aureus* | R | 10(15.9) | 12(19) | 15(23.8) | 30(47.6) | 12(19) | 31(49.2) | 7(11.3) | 24(39.1) | 54(85.7) | 59(93.7) |
|  | S | 53(84.1) | 51(81) | 48(76.2) | 33(52.4) | 51(81) | 32(50.8) | 55(88.7) | 39(61.9) | 9(14.3) | 4(6.3) |
| **Total** | R | 15(16.5) | 23(24.5) | 23(24) | 41(45) | 20(20.6) | 53(54.1) | 10(10.4) | 41(42.3) | 76(83.5) | 90(92.8) |
|  | S | 76(83.5) | 71(75.5) | 73(76) | 50(55) | 77(79.4) | 45(45.9) | 86(89.6) | 56(57.7) | 15(16.5) | 7(7.2) |

N: Number of tested strains; R: Resistant; S: Sensitive; Ptn: Pattern; FOX: Cefoxitin; GEN: Gentamicin; CIP: Ciprofloxacin; CHL: Chloramphenicol; SXT: Trimethoprim-Sulfamethoxazole; VAN: Vancomycin; ERY: Erythromycin; DA: Clindamycin; DOX: Doxycycline; PEN: Penicillin, NT: Not tested, GBS : Group B Streptococcus (*Streptococcus agalactiae*), CoNS: Coagulase negative staphylococci

### Antibiogram profile for Gram-negative isolates

Most of the GNB exhibited significantly high resistance to most of the tested antibiotics; for example, ampicillin (97.5%), ceftazidime (91.3%), ceftriaxone (91.3%) and aztreonam (90%), cefotaxime (83.8%), amoxicillin and clavulanic acid (77.6%) and cefoxitin (76.3%). Similarly, significant resistance level was also recorded for cefepime (75%), sulfamethoxazole-trimethoprim (71.3%), piperacillin-tazobactam (68.7%) and meropenem (56.3%). Low level resistance was recorded for amikacin (25%), ciprofloxacin (37.5%) and gentamicin (46.3%). *Acinetobacter* spp showed the highest resistance level to almost all tested antibiotics including penicillin, cephalosporins, and carbapenems and monobactam groups of antibiotics including: ampicillin (100%), aztreonam (100%), ceftazidime (100%), amoxicillin and clavulanic acid (100%), ceftriaxone (97.4%) and cefotaxime (92.3%). Low resistance level by *Acinetobacter* spp was recorded to amikacin (25%) (Table 5).

**Table 5:** Antimicrobial susceptibility pattern of Gram-negative bacteria at TASH, 2018

| Isolates | Ptn | Antimicrobial agent's n (%) |  |  |  |  |  |  |  |  |  |  |  |  |  |
| --- | --- | --- | --- | --- | --- | --- | --- | --- | --- | --- | --- | --- | --- | --- | --- |
|  |  | AMP | AZM | CTX | CRO | CTZ | FOX | FEP | AMC | TZP | MRP | AK | GEN | CIP | SXT |
| *Acinetobacter* spp | R | 39(100) | 39(100) | 36(92.3) | 38(97.4) | 39(100) | 37(94.8) | 34(87.2) | 32(100) | 34(87.2) | 29(74.4) | 14(25) | 27(69.2) | 18(46.1) | 31(79.5) |
|  | S | 0(0) | 0(0) | 3(7.7) | 1(2.6) | 0(0) | 2(5.2) | 5(12.8) | 5(0) | 5(12.8) | 10(25.6) | 25(75) | 12(30.8) | 21(53.9) | 8(20.5) |
| *E. coli* | R | 9(100) | 6(66.7) | 7(77.8) | 7(77.8) | 7(77.8) | 4(44.4) | 7(77.7) | 5(55.4) | 5(55.6) | 3(33.3) | 2(22.2) | 4(44.4) | 4(44.4) | 7(77.8) |
|  | S | 0(0) | 3(33.3) | 2(22.2) | 2(22.2) | 2(22.2) | 5(55.6) | 2(33.3) | 4(44.6) | 4(44.4) | 6(66.7) | 7(77.8) | 5(55.6) | 5(55.6) | 2(22.2) |
| *Enterobacter* spp | R | 2(100) | 2(100) | 2(100) | 2(100) | 2(100) | 2(100) | 2(100) | 2(100) | 2(100) | 1(50) | 0(0) | 1(50) | 2(100) | 2(100) |
|  | S | 0(0) | 0(0) | 0(0) | 0(0) | 0(0) | 0(0) | 0(0) | 0(0) | 0(0) | 1(50) | 2(100) | 1(50) | 0(0) | 0(0) |
| *K. oxytoca* | R | 4(100) | 3(75) | 3(75) | 4(100) | 3(75) | 3(75) | 3(75) | 1(50) | 2(50) | 1(25) | 1(25) | 1(25) | 2(50) | 2(50) |
|  | S | 0(0) | 1(25) | 1(25) | 0(0) | 1(25) | 1(25) | 1(25) | 1(50) | 2(50) | 3(75) | 3(75) | 3(75) | 2(50) | 2(50) |
| *K. pneumoniae* | R | 5(83.3) | 4(66.7) | 4(66.7) | 6(100) | 5(83.7) | 3(50) | 5(83.3) | 6(100) | 4(66.7) | 4(66.7) | 3(50) | 2(33.3) | 2(33.4) | 5(83.3) |
|  | S | 1(16.7) | 2(33.3) | 2(33.3) | 0(0) | 1(16.3) | 3(50) | 1(16.7) | 0(0) | 2(33.3) | 2(33.3) | 3(50) | 4(66.7) | 4(66.6) | 1(16.7) |
| *K. rhino* | R | 1(100) | 1(100) | 1(100) | 1(100) | 1(100) | 0(0) | 1(100) | 1(100) | 0(0) | 0(0) | 0(0) | 0(0) | 0(0) | 0(0) |
|  | S | 0(0) | 0(0) | 0(0) | 0(0) | 0(0) | 1(100) | 0(0) | 0(0) | 1(100) | 1(100) | 1(100) | 1(100) | 1(100) | 1(100) |
| *Pseudomonas* spp | R | 13(92.9) | 13(92.9) | 12(85.7) | 12(85.7) | 11(78.6) | 11(78.5) | 5(35.7) | 8(57.1) | 5(35.7) | 6(42.8) | 0(0) | 2(14.3) | 2(14.3) | 8(57.2) |
|  | S | 1(7.1) | 1(7.1) | 2(14.3) | 2(14.3) | 3(21.4) | 3(21.5) | 9(64.3) | 6(42.9) | 9(64.3) | 8(57.2) | 14(100) | 12(85.7) | 12(85.7) | 6(43.7) |
| *Serratia* spp | R | 4(100) | 3(75) | 1(25) | 2(50) | 4(100) | 1(25) | 3(75) | 3(75) | 2(50) | 1(25) | 0(0) | 0(0) | 0(0) | 1(25) |
|  | S | 0(0) | 1(25) | 3(75) | 2(50) | 0(0) | 3(75) | 1(25) | 1(25) | 2(50) | 3(75) | 4(100) | 4(100) | 4(100) | 3(75) |
| *Shigella* spp | R | 1(100) | 1(100) | 1(100) | 1(100) | 1(100) | 0(0) | 0(0) | 1(100) | 1(100) | 0(0) | 0(0) | 0(0) | 0(0) | 1(100) |
|  | S | 0(0) | 0(0) | 0(0) | 0(0) | 0(0) | 1(100) | 1(100) | 0(0) | 0(0) | 1(100) | 1(100) | 1(100) | 1(100) | 0(0) |
| Total | R | 78(97.5) | 72(90) | 67(83.8) | 73(91.3) | 73(91.3) | 61(76.3) | 60(75) | 59(77.6) | 55(68.7) | 45(56.3) | 20(25) | 37(46.3) | 30(37.5) | 57(71.3) |
|  | S | 2(2.5) | 2(10) | 13(16.2) | 7(8.7) | 7(8.7) | 19(23.7) | 20(25) | 17(22.4) | 25(31.3) | 35(43.7) | 60(75) | 43(53.7) | 50(62.5) | 23(28.7) |

N: Number of tested strains; R: resistant; S: Sensitive; %: percentage; Ptn: Pattern; AMP: ampicillin; AZT: Aztreonam; CTX : cefotaxime; CRO: ceftriaxone; CTZ: ceftazidime; FOX: Cefoxitin; FEP: Cefepime; AMC: Amoxicillin and clavulanic acid ; CHL: Chloramphenicol; MRP: Meropenem; AK: Amikacin; GEN: Gentamicin; CIP: ciprofloxacin; SXT: Sulfamethoxazole + trimethoprim

## Discussions

In the present study, out of 164 fomites and medical devices samples from swabs of normally clean hospital environments, 141(86%) were positive for bacterial contamination. Our result agreed with other reports where bacterial contamination was found to be very high such as report from Zimbabwe (86.2%) [14] and from Morocco (96.3%) [23]. In contrast to our result, lower bacterial contaminations were observed from studies conducted elsewhere; Gaza Strip (24.7%) [24], Sudan (29.7%) [25], Uganda (44.2%) [26], Nigeria (39.4%) [27] and Bahir Dar, Northwest Ethiopia (39.6%) [16]. Differences in hand hygiene, ventilation system, sterilisation and disinfection techniques could account for these discrepancies [1, 28, 29].

Higher levels of bacterial contamination observed in our study could be attributed primarily to the use of ineffective disinfectants during surface cleaning, and inadequate uses of standard precautions such as hand hygiene and contact precautions, as well as migration of the organisms through air flow or other means particularly in places where the ventlation system has not been not in place or not working properly [20]. Infrequent cleaning of inanimate surfaces and medical equipments could also contribute to poor microbial quality of the hospital surfaces [14, 30, 31]. This situation is prominently linked to hospitals which show unwillingness to put funds into contamination control such as the ventilation systems, those that lack information about the level of contamination and ineffectiveness of commonly used disinfectants in their hospital, and those with inappropriate waste controls.

The results of our study showed substantial contamination of hospital inanimate environments by varied groups of bacteria, including both Gram-positive (56.3%) and Gram-negative (43.7%). Comparable to our results, frequency of GPB from other studies in Ethiopia and abroad proved to be constituted the leading contaminating bacteria compared to GNB; for example, in Gondar, Ethiopia (60.5% vs 39.5%) [32], in Northwest, Ethiopia (81.6% vs 18.4%) [16], in Iran (60.7% vs 39.3%) [33] and in Nigeria (52.2% vs 47.8%) [34]. The dominance of GPB could be explained by the fact that these bacteria, being devoid of lipid-dominant desiccation prone outer membrane, have natural ability to retain their viability on abiotic hospital environments for several days to months [29, 33].

However, in contrast to our results, several authors from different countries reported that GNB were isolated more frequently than Gram-positive ones: for example, Zimbabwe (66.2% vs 33.82%) [14], Gaza Strip (51.6% vs 48.4%) [24] and Morocco (73.3% vs 26.7%) [23]. These variations may be due to different sampling times (e.g. during endemic vs outbreak situations), the presence of already colonized and/or infected patients during sampling, the use of different sampling techniques and culture methodologies, and variation in specific hospital sampling sites (e.g., OTs vs ICUs) [35–38]. In fact, in agreement to the latter reasoning, more GNB (67.4%; 62/92) than Gram-positive ones were obtained from ICUs inanimate environment even our finding.

Overall, *S. aureus* was the most frequently isolated bacteria (39.8%) followed by *Acinetobacter* spp (18.9%) and CONS (15.5%). *S. aureus* and CONS were also the most frequently isolated bacteria from previous other studies such as Ethiopia [39], Nigeria [34] and Zaria, Nigeria [27]. *S. aureus* constitute part of the normal human flora, inhabiting the skin, mucous membranes [40] and regularly shed onto the hospital environment by patients and medical personnel, whereupon they persist [14]. This isolates were also considered as the potential pathogenic bacteria that result in nosocomial infections and indicators of inadequate clinical surface hygiene in hospital environments [17, 25, 41]. Moreover, these bacteria were also resistant to common disinfectant methods and hence spread easily in the environment, which enables them to colonize and infect the patients receiving health care service at the facility [24, 33].

Among the different hospital environments and hospital items examined, the highest bacterial contaminated samples were taken from bed linens, environmental surface and beds, similar to the observations from other studies in Ethiopia and abroad [3, 27, 33, 36]. Bed linens and bed were mainly contaminated by *Acinetobacter* spp (20.5% and 15.4%), CONS (7.1% and 14.3%), and *S. aureus* (12.7% and 7.9%), respectively. Comparable results were obtained on beds and linens samples from studies conducted in Iran [33] and Nigeria [27]. The sources of such contaminations could be cross-contamination from a patient’s flora, health care workers’ hands, contaminated storage carts, or due to contamination during the washing process especially that of bed linens [33, 35, 37].

In our study, sinks were mainly colonized by *S. aureus* (7.7%, 6/63), *Pseudomonas* spp (7.1%, 1/14) and *Acinetobacter* spp (5.1%, 2/39), which is in line with several reports that hospital associated outbreaks in critical care wards occur largely due to the opportunistic pathogen [14, 27, 42]. This could be linked to the fact that the moist hospital environments, particularly sinks, are conducive for persistence of these bacteria, which are known to have the ability to form biofilms in water, sinks, toilets, showers and drains [43, 44]. Moreover, acquisition of multiple virulence determinants and intrinsic resistance to commonly used antibiotics and disinfectants by these pathogens may result in maintaining their viability and hence persistence under such harsh environments [43, 45].

Bloodstream infection and ventilator-associated pneumonia especially in the intensive care units are usually linked to device contamination such as central venous catheters, urinary catheters and ventilators [46]. In our study, ventilators were frequently contaminated by *Klebsiella* spp (27.3%, 3/11), which was also reported from a study conducted in Iran (54.4%, 6/11) [33]. Source of contamination of ventilators by *K. pneumoniae* might be from the aspiration of secretions from the oropharynx of colonized patients, where staff hands may act as the transmission vehicle [47, 48].

In regards to antimicrobial resistance profile of the isolates, our results showed high proportions of drug resistance, where most of the GNB were highly resistant to most of the tested antibiotics such as ampicillin (97.5%), ceftazidime (91.3%), ceftriaxone (91.3%), aztreonam (90%), cefotaxime (83.8%), cefoxitin (76.3%), and amoxicillin and clavulanic acid (77.6%), which is in line with similar resistance rates from other studies conducted elsewhere like Gaza in Palestine [24], Morocco [3] and Sudan [25]. Increased resistance to β-lactams antibiotics is due to the selective pressure exerted by the antibiotics [49]. Because these tested antimicrobials represent the antibiotics most frequently used in practice, serious problems can be encountered while prescribing those antibiotics [3]. One way of fighting such a rise of resistance should include establishing guidelines for prescribing antibiotics [16] based on locally generated antimicrobial resistance data such as the findings from this study.

On the other hand, low resistance level was recorded to non-beta-lactam antimicrobials such as, amikacin (25%) and ciprofloxacin (37.5%). Comparable results were recorded from studies conducted from Sudan for amikacin (23.5%) and ciprofloxacin (42.7%) [25]. Still lower resistance rate was documented for these two antibiotics in Palestine for amikacin (6.1%) and ciprofloxacin (27.3%) [24], possibly an area where they may not routinely be prescribed for community and/or hospital acquired infections.

Not surprisingly, GPB demonstrated elevated resistance to penicillin (92.8%), cefoxitin (83.5%) and erythromycin (54.1%). Similarly, high resistance level was also reported from Ethiopia by a Meta-analysis study for penicillin and erythromycin with a pooled resistance level of 99.1% and 97.2%, respectively [50]. Moreover, similar resistance level was also reported from Uganda for penicillin (93%) [26]. Of the 64 *S. aureus* isolates obtained in this study, 54 (85.7%) were MRSA, which is close to the rate reported from Zimbabwe (100%) [14], although much higher than the rate from Uganda (52%) [51].

In this study, vancomycin resistance was demonstrated by 12 (19%) *S. aureus* (VRSA), 5(17.9%) CONS and 1(33.3%) *Enterococcus* spp. Vancomycin resistant *Staphylococci* were also reported in a study from Zimbabwe, where 40% of *S. aureus* and 23.5% of CONS were vancomycin resistant, despite its scarcity in usage [14]. It has been suggested that patients at risk for VRSA are co-infected or co-colonized with VRE and MRSA, which enables conjugative transfer of vanA gene from VRE to MRSA in a biofilm environment leading to a VRSA strain [33, 52].

## Conclusions

In this study, bacterial samples were sought for and isolated only from the environmental surfaces; not from patients and hands of health professionals. *S. aureus*, *Acinetobacter* spp and CONS form the majority of the environmental contaminants most likely to cause HAIs. We concluded that special attention to infection control policies, antimicrobial resistance screening, good clinical practice and cleaning techniques are needed to reduce the potential risk of pathogenic bacteria and resistant strain transmission among hospital staff and patients. Our results may be indicative evidence that bacterial environmental contamination is possibly contributing to HAIs and MDR strain dissemination in the hospital environment and further large scale investigations are needed.

## Additional files

**S 1 Table**: Morphological and biochemical characterization of gram-positive bacteria isolated from environmental samples at Tikur Anbessa Specialized Hospital, Ethiopia, 2018 (DOC 35 kb).

**S 2 Table**: Morphological and biochemical characterization of gram-negative bacteria isolated from environmental samples at Tikur Anbessa Specialized Hospital, Ethiopia, 2018 (DOC 35 kb).

**S 3 Table**: Data description (DOC 17 kb).

## Acknowledgements

The authors here by thank Addis Ababa University (AAU) and Armauer Hansen Research Institute (AHRI) for their financial and material support and Tikur Anbessa Specialized Hospital staff.

## Data Availability

The dataset supporting the findings of this article have been attached as supplementary information files.

